# Contrasting dynamics of soil fungal functional groups in the plant rhizosphere

**DOI:** 10.1101/2024.08.30.610541

**Authors:** Na Wei, Madelynn Nakaji-Conley

## Abstract

**Background and aims:** Soil microbiomes, critical for plant productivity and ecosystem functioning, mediate essential functions such as pathogenesis, mutualism, and decomposition through different fungal functional groups. Yet, our understanding of the dynamics of co-existing soil fungal functional groups in the plant rhizosphere remains limited.

**Methods:** By leveraging a ‘natural’ experiment in urban farming with fields of different ages and multiple plant genotypes, we tracked the relative abundance, richness, and microbial networks of plant pathogens, mycorrhizal fungi, and saprotrophic fungi across fields over two years.

**Results:** We observed an increase in the relative abundance of plant pathogens in older fields relative to younger fields, supporting the prediction of pathogen accumulation over time. In contrast, there was a decrease in the relative abundance of mycorrhizal fungi in older fields. Unlike plant pathogens and mycorrhizal fungi, the relative abundance of saprotrophic fungi remained similar among fields. While the richness of plant pathogens and saprotrophic fungi were similar across fields, the community structure of both groups differed between younger and older fields. For mycorrhizal fungi, the richness declined in older fields and over the two years. These dynamics led to distinct microbial networks, with decreased network links for mycorrhizal fungi and increased links for saprotrophic fungi in older fields, whereas the links for plant pathogens remained similar across fields.

**Conclusion:** Our study reveals contrasting dynamics of essential soil fungal functional groups in the plant rhizosphere, and provides a predictive insight into the potential shifts in soil function and their impact on plant productivity.

## Introduction

Soil microbiomes are key to plant productivity and ecosystem functioning (Gilbert 2002; Smith and Smith 2011; Fierer 2017; Tedersoo et al. 2020; Lebreton et al. 2021; Liu et al. 2022). The essential functions of soil microbiomes, such as pathogenesis, mutualism, and decomposition, are mediated by different functional groups. For example, soil-borne fungal pathogens have the potential to induce plant pathogenesis (Gilbert 2002; Madden and Wheelis 2003; Möller and Stukenbrock 2017). The accumulation and diversity of pathogens influence plant growth and vegetation dynamics and are negatively linked to the resilience and resistance of plant productivity (Semchenko et al. 2018; Liu et al. 2022; Kohout et al. 2024). On the other hand, mycorrhizal fungi, including both arbuscular mycorrhizae and ectomycorrhizae, offer protection to plants, by enhancing nutrient and water access and protecting against pathogens and stress (Tedersoo et al. 2020; Chandrasekaran and Paramasivan 2022; Marro et al. 2022; Qiu et al. 2022; Liu et al. 2024). Mycorrhizal fungi can also impact plant–plant interactions, mediating species coexistence and diversity maintenance through belowground mycorrhizal networks (Tedersoo et al. 2020). Saprotrophic fungi, as decomposers, play a vital role in breaking down complex organic matter and releasing essential nutrients, such as nitrogen, phosphorus, and other minerals back into the soil, thereby facilitating carbon and nutrient cycling (Hättenschwiler et al. 2005; Grinhut et al. 2007; Lebreton et al. 2021). The diversity of saprotrophic fungi has been found to be positively linked to the stability, resilience, and resistance of plant productivity (Liu et al. 2022). Despite their significance, our knowledge of the dynamics of different fungal functional groups that co-exist in rhizosphere soil microbiomes is limited (Zanne et al. 2020; Martinović et al. 2021).

Soil fungal functional groups can be dynamic in the plant rhizosphere. For example, fungal pathogens can accumulate over time, especially in monocultures or ecosystems with low plant species diversity (Cook 2006; Li et al. 2014; Peralta et al. 2018; Wang et al. 2023a). The buildup of specialist pathogens can occur in environments where host plants are continuously or repeatedly present. In contrast, the pathogen dilution theory predicts that in ecosystems with high plant diversity, this diversity can buffer against pathogen accumulation (Keesing et al. 2006). This can occur through mechanisms such as host regulation, where interspecific competition reduces the abundance of compatible host plants, and through encounter reduction, where non-host plants reduce pathogen transmission (Keesing et al. 2006). Pathogen accumulation over time can also occur through plant–soil feedbacks, where plant species contribute to the proliferation of host-specific pathogens that in turn affect the host plants (van der Putten et al. 2013; Cesarano et al. 2017; Semchenko et al. 2018; Bennett and Klironomos 2019). These soil legacy effects point to the long-lasting accumulation and impact of soil-borne pathogens over time, which can persist even after the host plants are no longer present (van der Putten et al. 2013; Heinen et al. 2020). In the case of accumulated pathogens, mycorrhizal fungi may decrease due to, for example, reduced host resource allocation to mutualistic fungi as a result of weakened host health (Tedersoo et al. 2020). Moreover, the abundance of mycorrhizal fungi can be dynamic contingent upon resource availability, as high resource availability reduces plant demand for and investment in mutualistic symbiosis (Treseder 2004; Werner and Kiers 2015; Lekberg et al. 2021). Saprotrophic fungi, on the other hand, may increase if there is an accumulated availability of organic matter in the soil through litterfall and root turnover, providing substrates for decomposition and influencing saprotroph activities (Hättenschwiler et al. 2005; Grinhut et al. 2007). The dynamics of different fungal functional groups are, nevertheless, not independent (Bödeker et al. 2016; Bennett and Klironomos 2019; Liu et al. 2024). The synergistic and antagonistic interactions (Raaijmakers et al. 2008) can lead to complex dynamics among different fungal functional groups over time, which are essential for plant productivity but remain poorly understood in both natural and managed ecosystems.

Plant species and genotypes can affect microbial community assembly in both wild and cultivated plants (Wei and Ashman 2018; Wei et al. 2021; Wei et al. 2022; Wei and Tan 2023), by impacting various aspects such as plant immune system, metabolic pathway, physiology, and morphology (Shakir et al. 2021). For example, plant genetic factors that influence immunity responses can modulate different levels of tolerance and resistance to fungal pathogens among different genotypes (Ferreira et al. 2007; Kushalappa et al. 2016; Teixeira et al. 2019). Plant genotypes may also vary in partner quality for and partner selection on mycorrhizal fungi, due to differential abilities in carbon investment, discrimination against less beneficial mycorrhizal partners, or root morphology such as coarseness, thereby influencing mycorrhizal association (Werner and Kiers 2015). Furthermore, genotypes that vary in the quantities and qualities of plant litter (e.g., chemical composition, nutrient content, physical structure and texture) and thereby affect saprotroph populations (Schweitzer et al. 2005; Semchenko et al. 2021). Nevertheless, our knowledge of the dynamics of these fungal functional groups across different plant genotypes remains scarce.

To understand the dynamics of soil fungal functional groups, specifically plant pathogens, mycorrhizal fungi, and saprotrophic fungi in the rhizosphere, we leveraged a ‘natural’ experiment in urban farming that involved multiple genotypes and fields of different ages. By tracking the dynamics of soil fungal functional groups across fields over two years, we aimed to test the following hypotheses (Fig. 1): (1) the relative abundance and richness of plant pathogens increase in older fields relative to younger fields and show an increase over the two years, (2) mycorrhizal fungi decrease in older fields and over time, and (3) saprotrophic fungi increase in older fields and over time. Additionally, we hypothesize that (4) these dynamics would lead to changes in microbial networks across fields and over time. Lastly, we predict that (5) the dynamics of plant pathogens, mycorrhizal fungi, and saprotrophic fungi may vary among plant genotypes.

**Fig. 1.**
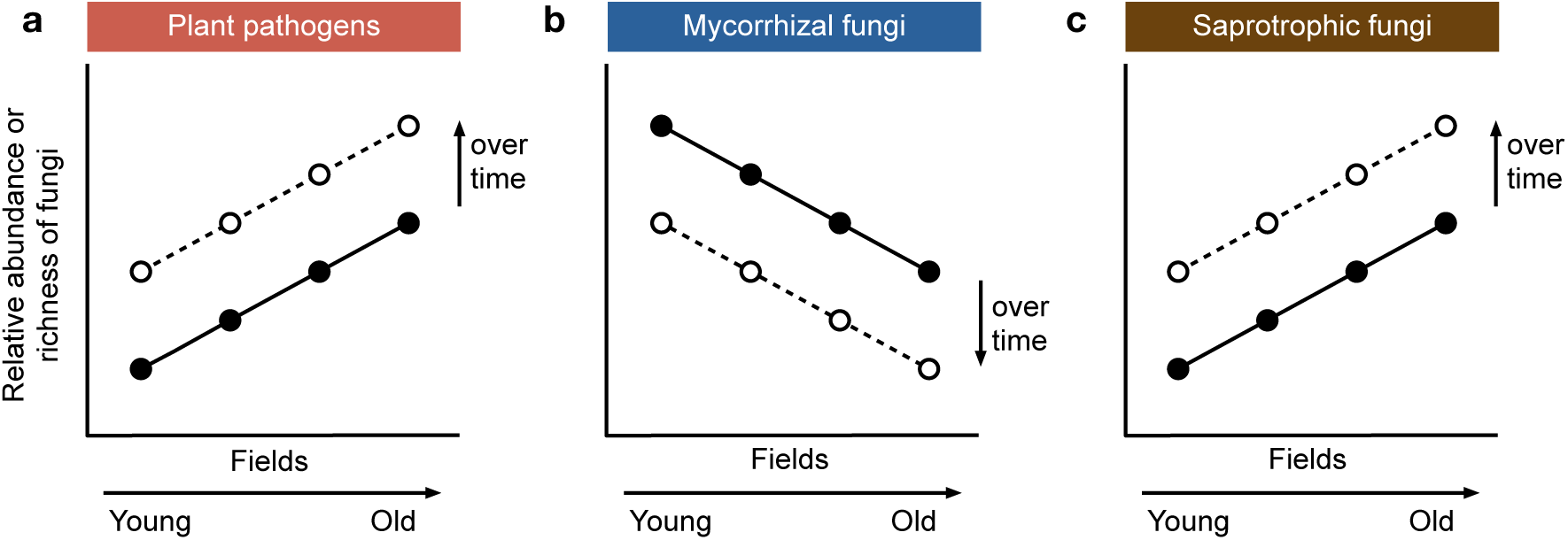
Hypotheses of changes in soil fungal functional groups among fields of different ages. (a) The relative abundance and richness of plant pathogens are predicted to increase in older fields relative to younger fields. (b) Mycorrhizal fungi are predicted to decrease in older fields. (c) Saprotrophic fungi are predicted to increase in older fields. These patterns (a–c) are expected to be more pronounced over time (solid lines vs. dashed lines).

## Materials and methods

### Study site and soil collection

We conducted the study at an urban farm in Chesterland, Ohio (41.554652°N, 81.324045°W). Four perennial strawberry fields, denoted as field A (planted in 2019), field B (2018), field C (2017), and field D (2016), were selected for investigation (Table S1). Each field (*c*. 50 m × 100 m) consisted of 35–55 matted rows growing 3–6 strawberry genotypes (*Fragaria* ×*ananassa* “AC Valley Sunset”, “Allstar”, “DaRoyal”, “DarSelect”, “Earliglow”, and “Wendy”; Table S1). Within each field, we collected soil microbiome samples from the plant rhizosphere in three randomly selected plots (20 cm × 20 cm) per genotype across non-adjacent rows during June 14– 15, 2021. We repeated the sampling at the same locations on June 21, 2022, with the exception of field B, which was removed by the farm in 2022. Specifically, for soil microbiome collection, we took soil from the root zone of a strawberry plant using an ethanol-cleaned stainless steel hand transplanter. Each sample, comprising 0.5 mL of soil, was transferred into a sterile 2 mL microcentrifuge tube. The samples (*N* = 96) were stored at -20 °C within 3 h after collection.

### Soil microbiome DNA extraction and sequencing

Soil microbial DNA was extracted using cetyltrimethylammonium bromide (CTAB) and purified using polyethylene glycol (PEG) 8000, following the previous protocol (Wei and Tan 2023). Briefly, soil samples were lysed with 1 mL sterile CTAB buffer (2% w/v CTAB, 100 mM Tris-HCl, 20 mM EDTA, 1.4 M NaCl, 5 mM ascorbic acid, and 10 mM dithiothreitol) on a Vortex Genie 2 (Scientific Industries, Bohemia, New York) for 40 min. After brief centrifuging, the supernatant was transferred to a new sterile 2 mL microcentrifuge tube, and an equal volume of chloroform : isoamyl alcohol (24 : 1) was added for phase separation at 13,200 rpm for 5 min. DNA was then recovered by adding the upper phase to two volumes of cold pure ethanol overnight at -20 °C. Pelleted DNA was washed with 500 µL of cold 70% ethanol and eluted in sterile TE buffer. The DNA was further purified with an additional round of chloroform : isoamyl alcohol phase separation, and was then recovered by adding the upper phase to an equal volume of sterile PEG 8000 (20% w/v PEG 8000, 2.5 M NaCl) and incubating at 37 °C for 30 min. Pelleted DNA was washed with cold 70% ethanol and eluted in 60 µL sterile TE buffer. The DNA samples were sent to the Argonne National Laboratory for fungal library preparation (ITS1f–ITS2) and sequencing using Illumina MiSeq (paired-end 250 bp).

### Fungal sequence analysis

Demultiplexed paired-end (PE) reads were used for identifying fungal amplicon sequence variants (ASVs) using package DADA2 v1.20.0 (Callahan et al. 2016) in R v4.2.2 (R Core Team 2022). Following the previous pipeline (Wei et al. 2021; Wei et al. 2022), the PE reads were screened to remove potential primer contaminations and quality filtered [minLen = 50, maxN = 0, truncQ = 2, maxEE = c(2, 2)], prior to specific variant identification that took into account sequence errors. The PE reads were then end joined (minOverlap = 20, maxMismatch = 0) for ASV detection. After chimera removal, fungal ASVs were assigned with taxonomic identification based on the UNITE reference database (v8.0 dynamic release) using QIIME 2 v2023.2 (Bolyen et al. 2019). The fungal data were further filtered. First, we removed non-fungal ASVs. Second, we filtered out samples of low reads (<1000 reads; *N* = 1). Third, we conducted rarefaction analysis using the package iNEXT (Cao et al. 2018) to confirm that the sequencing effort was sufficient to capture the fungal richness and to guide reads normalization (Fig. S1). Based on the rarefaction result, we normalized per-sample reads to the same number (8,000) across samples, following the previous work (Wei et al. 2021; Wei et al. 2022). Lastly, we removed low-frequency ASVs (<0.001% of total observations). The final fungal data consisted of 2239 ASVs across the remaining 95 samples. Fungal functional groups (potential plant pathogens, mycorrhizal fungi including both arbuscular and ectomycorrhizal fungi, and saprotrophic fungi; Table S2) were identified using FungalTraits v1.2 (Põlme et al. 2021).

### Statistical analyses

To evaluate whether older fields harbor more plant pathogens, fewer mycorrhizal fungi, and more saprotrophic fungi (Fig. 1), we conducted general linear models (LMs) with field and genotype as the predictors. The response variable was the relative abundance or richness of individual fungal functional groups. Due to data imbalance resulting from the removal of field B in 2022, LMs were conducted for 2021 and 2022 separately. To evaluate whether plant pathogens increased, mycorrhizal fungi decreased, and saprotrophic fungi increased over the two years (2021 vs. 2022), we conducted such comparisons for each field using LMs (predictors: collection year, genotype, and their interactions). Likewise, to evaluate whether and how fungal functional groups changed over the two years for each genotype, we conducted such comparisons for each genotype using LMs (predictors: collection year, field, and their interactions). Statistical significance (type III sums of squares), least-squares means (LS-means) of predictors, and the planned contrasts within the above LMs were assessed using package emmeans (Lenth et al. 2019).

To evaluate how the community structure of fungal functional groups varied among fields each year, we conducted permutational multivariate analyses of variance (PERMANOVA) based on Bray–Curtis dissimilarity using the package vegan (Oksanen et al. 2019). The predictors included field and genotype. We visualized the communities using principal coordinates analyses (PCoA).

To evaluate the potential interactions within and between fungal functional groups, we constructed microbial correlational networks based on Kendall’s correlation (coefficient ≥0.6 or ≤-0.6), a non-parametric method with greater robustness than Spearman’s correlation, using the package correlation (Makowski et al. 2020). Networks were built separately for each field and year based on the ASVs of plant pathogens, mycorrhizal fungi, and saprotrophic fungi using the package igraph (Csárdi et al. 2024). Link-level information was calculated within functional groups (pathogen-pathogen, mycorrhiza-mycorrhiza, and saprotroph-saprotroph) and between functional groups (pathogen-mycorrhiza, pathogen-saprotroph, and mycorrhiza-saprotroph).

## Results

### Changes in the relative abundance and richness of fungal functional groups among fields

Consistent with the prediction that older fields harbor more plant pathogens (Fig. 1a), the relative abundance of plant pathogens was lowest in field A that was planted in 2019 and highest in field C that was planted in 2017 (sampling year 2021; LS-mean, field A = 0.08 ± 0.03; field C = 0.16 ± 0.02; LS-mean contrast, *F* = 4.5, df = 1, *P* = 0.039; Fig. 2a). Plant pathogens increased from 2021 to 2022 in the younger field (field A, 2021 vs. 2022; LS-mean contrast, *F* = 5.3, df = 1, *P* = 0.040) and remained relatively unchanged in the older fields (field C and field D; both *P* > 0.05; Fig. 2a). Consistent with the prediction that older fields harbor fewer mycorrhizal fungi (Fig. 1b), the relative abundance of mycorrhizal fungi was highest in field A (planted in 2019) and lowest in field D (planted in 2016) in 2021 (LS-mean, field A = 0.15 ± 0.03; field D = 0.03 ± 0.03; LS-mean contrast, *F* = 8.3, df = 1, *P* = 0.006; Fig. 2b). From 2021 to 2022, mycorrhizal fungi decreased dramatically and consistently across fields (LS-mean contrast, field A, *F* = 45.0, df = 1, *P* < 0.001; field C, *F* = 9.1, df = 1, *P* = 0.006; field D, *F* = 5.3, df = 1, *P* = 0.033; Fig. 2b). Different from plant pathogens and mycorrhizal fungi, saprotrophic fungi remained similar among fields in 2021 and 2022 (LS-mean contrast, all *P* > 0.05; Fig. 2c).

**Fig. 2.**
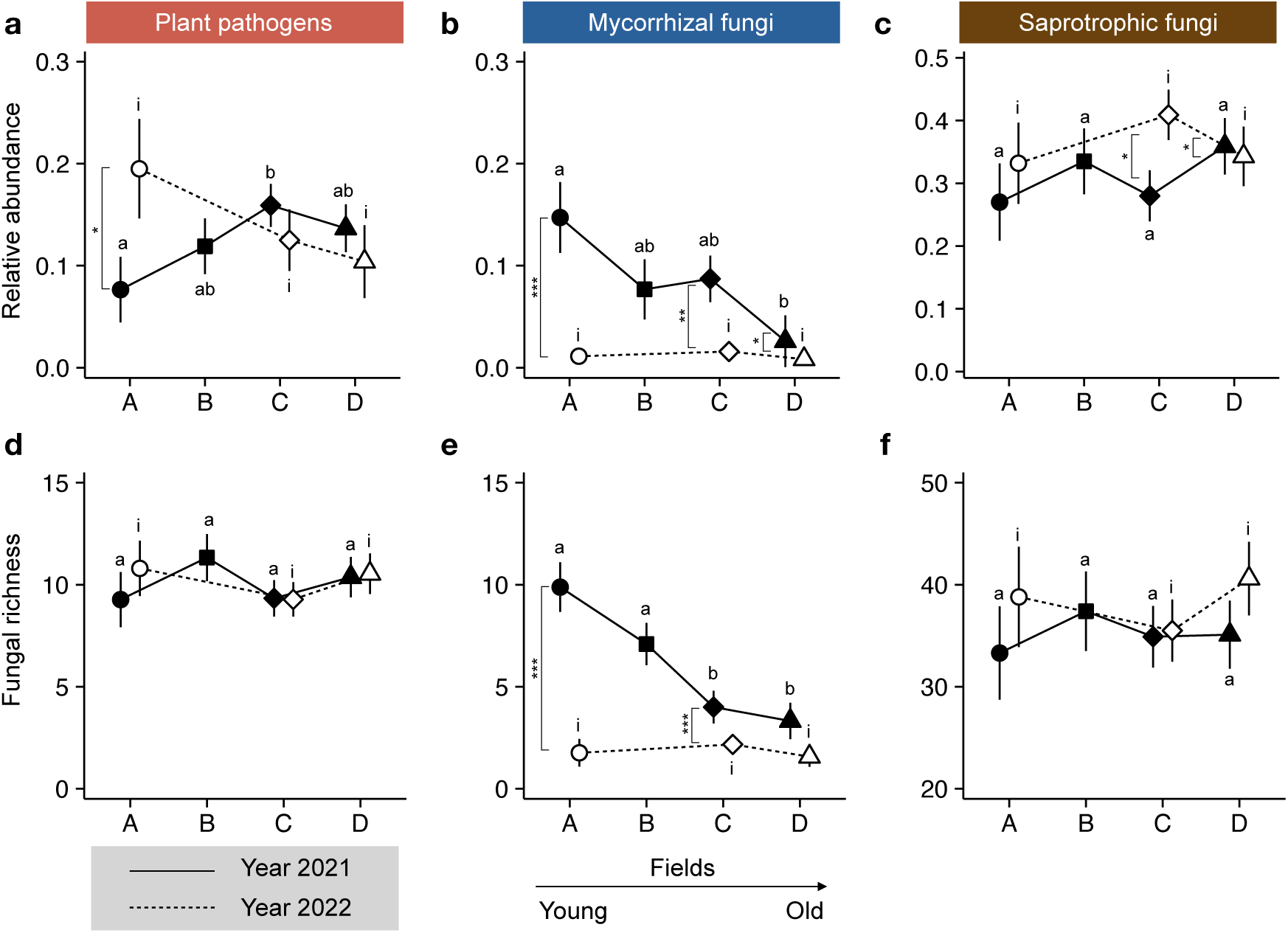
The relative abundance and richness of soil fungal functional groups vary among fields of different ages. The least-squares means (LS-means) of the relative abundance (a-c) and richness (d-f) of plant pathogens, mycorrhizal fungi, and saprotrophic fungi are plotted with error bars (1 SE) across the four fields: field A (planted in 2019), field B (2018), field C (2017) and field D (2016), after controlling for the effect of genotype in general linear models (LMs) for the years 2021 (solid symbols) and 2022 (open symbols). Field B was removed by the farm in 2022. LS-mean contrasts among fields are indicated by letters (a and b for 2021; i for 2022), and LS-mean contrasts between 2021 and 2022 are shown vertically. \*\*\**P* < 0.001; \*\**P* < 0.01; \**P* < 0.05.

Different from the relative abundance, the richness of plant pathogens remained similar among fields and over the two years (LS-mean contrast, all *P* > 0.05; Fig. 2d). In contrast, for mycorrhizal fungi (Fig. 2e), consistent with the expectation that older fields harbor fewer mycorrhizal fungi (Fig. 1b), the richness was lower in the older fields (C and D) than that in the younger fields (A and B) in 2021 (Fig. 2e). From 2021 to 2022, the richness of mycorrhizal fungi decreased dramatically (LS-mean contrast, field A, *F* = 58.1, df = 1, *P* < 0.001; field C, *F* = 21.7, df = 1, *P* < 0.001; Fig. 2e). For saprotrophic fungi, the richness remained similar among fields and over the two years (LS-mean contrast, all *P* > 0.05; Fig. 2f), similar to their relative abundances (Fig. 2c).

### Changes in the community structure of fungal functional groups among fields

The communities of plant pathogens in the younger fields differed from those in the older fields in 2021 (PERMANOVA, field A vs. C, 13% of total variation, *F* = 3.7, *P* = 0.003; field B vs. C, 14%, *F* = 4.7, *P* = 0.001; Fig. 3a) and 2022 (field A vs. C, 9%, *F* = 2.7, *P* = 0.001; field A vs. D, 10%, *F* = 2.6, *P* = 0.010; Fig. 3a). Such differences in the compositional variation of plant pathogens were primarily caused by higher *Olpidiaster* in field C and higher *Pilidium* from 2021 to 2022 in field A, while the dominant plant pathogens across fields belonged to *Fusarium* (Fig. 4a). The communities of mycorrhizal fungi that were dominated by *Rhizophagus* (Fig. 4b) were similar across fields in both years (Fig. 3b). Yet, there was an increased loss of mycorrhizal associations in the older fields (Fig. 4b), consistent with the decreased relative abundance and richness of mycorrhizal fungi in the older fields (Fig. 2). For saprotrophic fungi, while the relative abundance and richness were similar across fields (Fig. 2), the communities in the younger fields differed from those in the older fields in 2021 (field A vs. C, 5%, *F* = 1.3, *P* = 0.051; field A vs. D, 7%, *F* = 1.7, *P* = 0.011; field B vs. C, 10%, *F* = 3.1, *P* = 0.001; field B vs. D, 7%, *F* = 1.9, *P* = 0.007; Fig. 3c) and 2022 (field A vs. C, 8%, *F* = 2.3, *P* = 0.003; field A vs. D, 9%, *F* = 2.3, *P* = 0.001). The communities of saprotrophic fungi were dominated by diverse classes including, for example, *Leotiomycetes*, *Agaricomycetes*, *Sordariomycetes*, and *Dothideomycetes* (Fig. 4c).

**Fig. 3.**
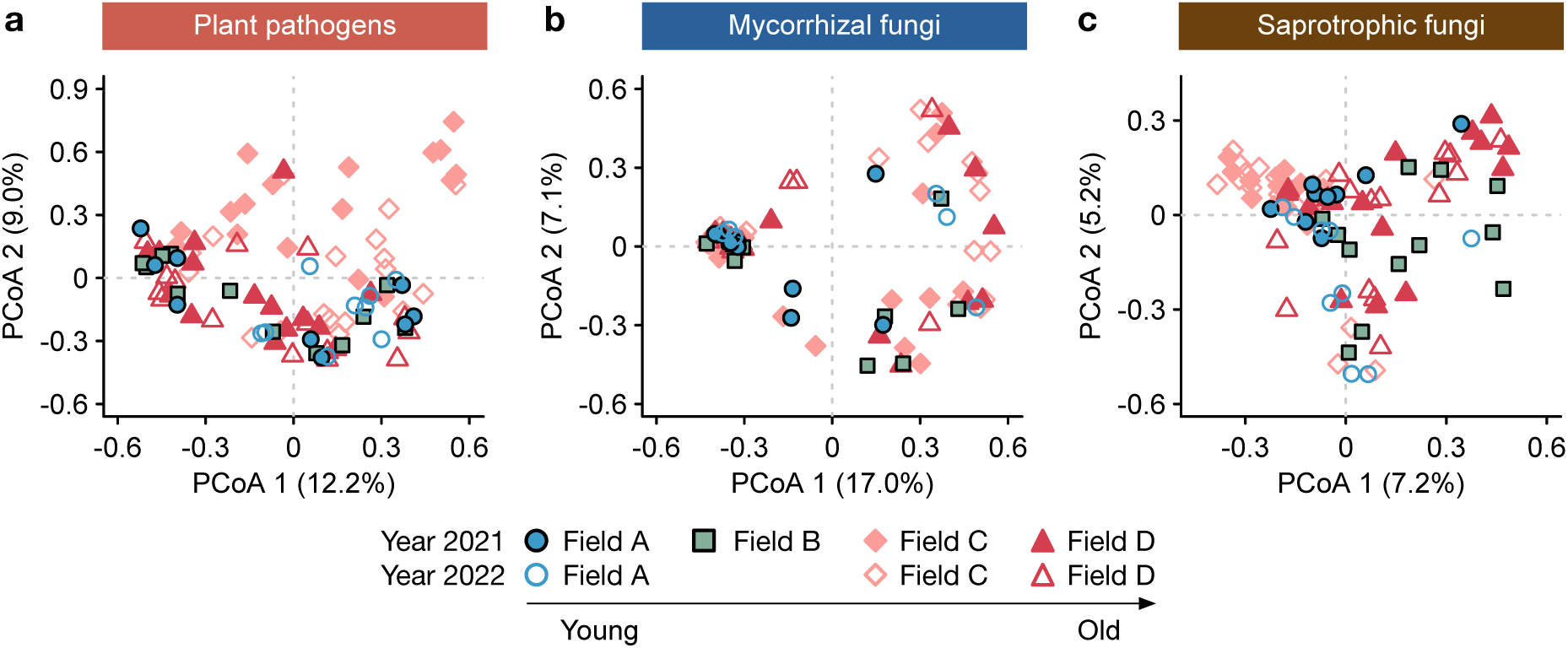
The communities of soil fungal functional groups vary among fields of different ages. (a) Plant pathogens showed significant compositional variation among fields in both 2021 and 2022, with younger fields (A and B) differing from the older fields (C and D), as revealed by principal coordinates analyses (PCoAs). (b) Mycorrhizal fungi did not differ significantly among fields in 2021 and 2022. (c) Saprotrophic fungi in younger fields (A and B) differed significantly from those in older fields (C and D) in 2021 and 2022. Field B was removed by the farm in 2022.

**Fig. 4.**
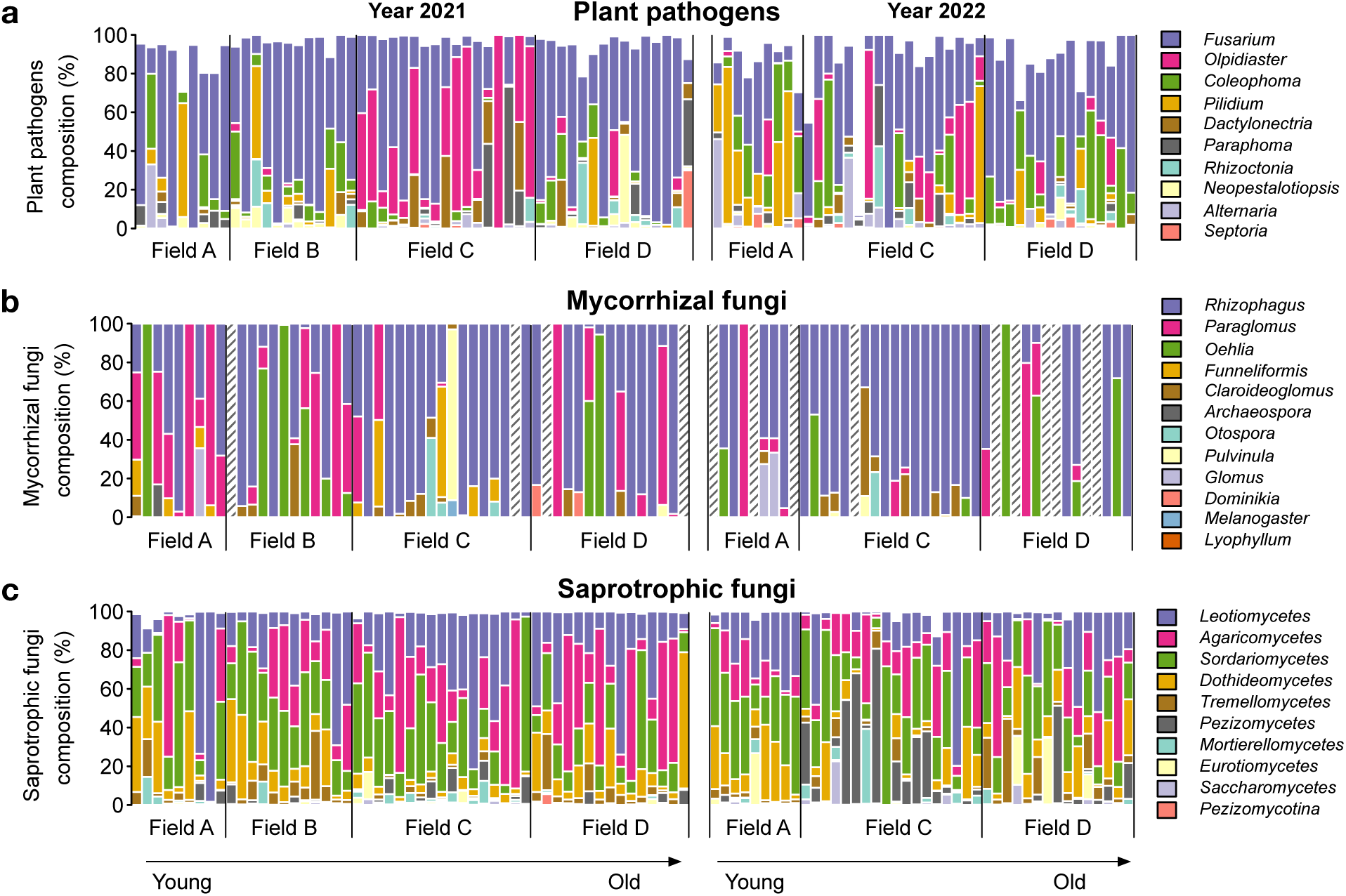
The composition of major taxonomical ranks of soil fungal functional groups. (a) The top 10 most abundant genera of plant pathogens, (b) all 12 genera of mycorrhizal fungi, and (c) the top 10 most abundant classes of saprotrophic fungi are shown across fields of different ages in 2021 and 2022. Hatched bars indicate the absence of mycorrhizal fungi, especially in older fields and in 2022. Field B was removed by the farm in 2022.

### Variation in microbial networks among fields

Microbial networks were dominated by positive connections (or links) both within and between fungal functional groups (Fig. 5). The negative links occurred primarily in the younger fields (field A and B; Fig. 5a,b,e). The proportion of mycorrhiza-mycorrhiza links was lower in the older fields compared to younger fields in 2021, and it consistently decreased across fields from 2021 to 2022 (Fig. 5). Such changes were due to a reduction in mycorrhizal fungi in the older fields and over the two years (Fig. 2). In 2021, mycorrhizal fungi accounted for 25% and 19% of the network nodes in the younger fields (A and B, respectively) but 13% and 12% in the older fields (C and D, respectively). In 2022, mycorrhizal nodes dropped to 5–7% across all fields (Fig. 5). In contrast, the proportion of saprotroph-saprotroph links increased from younger to older fields in 2021 and continued to rise from 2021 to 2022 (Fig. 5), which was due to the increased proportion of network nodes (2021: 61% and 65% in field A and B, respectively, and 73% and 72% in C and D, respectively; 2022: 78–80% across fields). The proportion of pathogen-pathogen links was relatively stable across fields in both years (Fig. 5), because the richness of plant pathogens (Fig. 2) and their network node proportions were similar across fields (2021: 14–15% across fields; 2022: 13–17%).

**Fig. 5.**
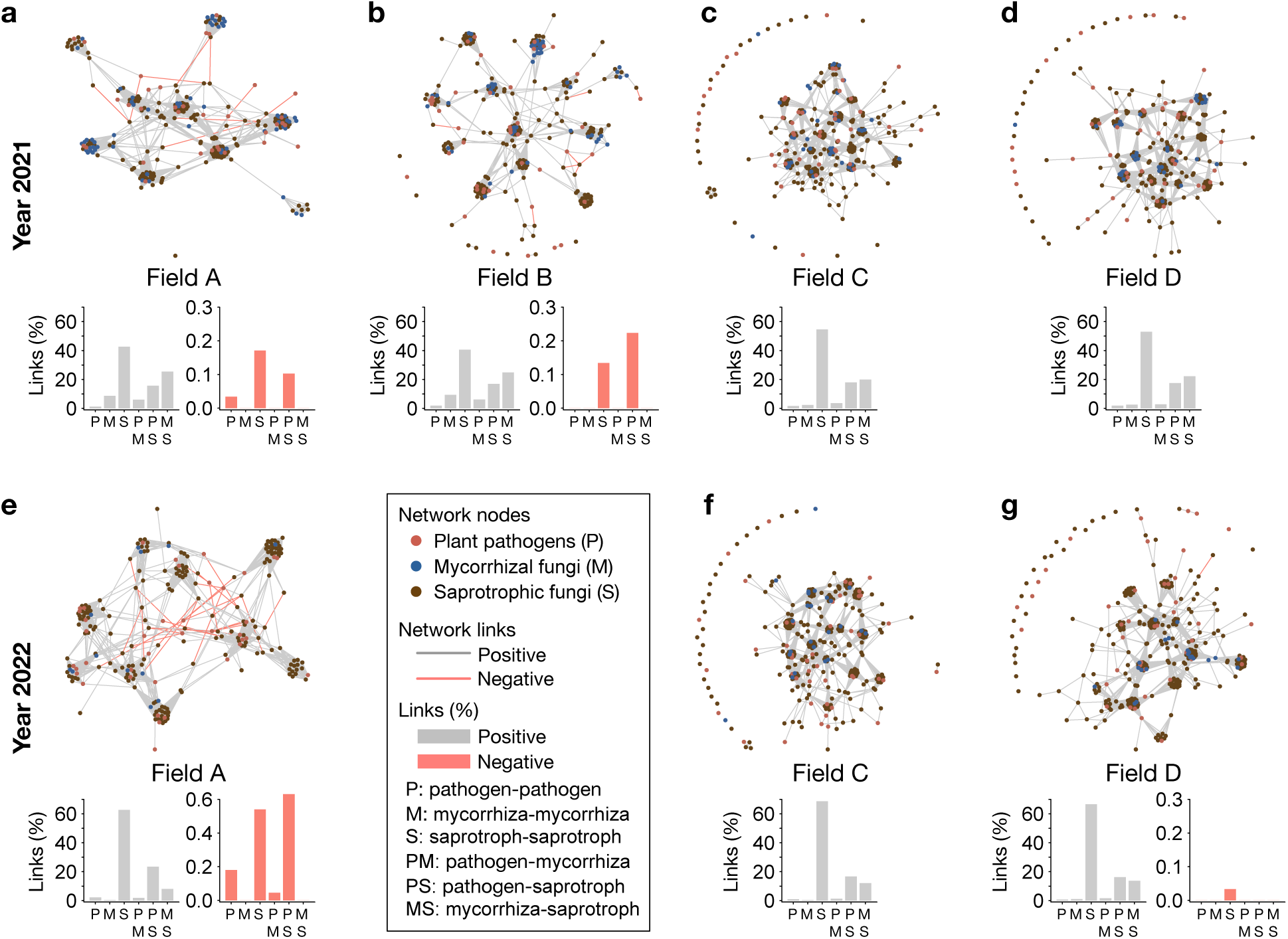
Microbial correlation networks of soil fungal functional groups across fields of different ages. Networks were constructed based on Kendall’s correlation coefficients (≥ 0.6 or ≤ -0.6) for plant pathogens (dark pink nodes), mycorrhizal fungi (blue nodes), and saprotrophic fungi (brown nodes) among samples in each field for the years 2021 (a-d) and 2022 (e-g). Positive correlations are indicated by grey links, while negative correlations are shown by salmon links within the networks. The percentage of microbial links (%) within and between functional groups are summarized by the bar plots. Field B was removed by the farm in 2022.

### Similar fungal functional groups among genotypes

The relative abundance and richness of plant pathogens and saprotrophic fungi were similar among genotypes in both years (Fig. 6a,c,d,f). For mycorrhizal fungi, the relative abundance and richness decreased from 2021 to 2022 in most genotypes: Allstar (LS-mean contrast, relative abundance, *F* = 12.8, df = 1, *P* = 0.004, Fig. 6b; richness, *F* = 48.7, df = 1, *P* < 0.001, Fig. 6e), DaRoyal (relative abundance, *F* = 7.4, df = 1, *P* = 0.027; richness, *F* = 7.0, df = 1, *P* = 0.029), DarSelect (relative abundance, *F* = 11.2, df = 1, *P* = 0.010; richness, *F* = 10.7, df = 1, *P* = 0.011), and Earliglow (relative abundance, *F* = 7.1, df = 1, *P* = 0.056; richness, *F* = 25, df = 1, *P* = 0.007).

**Fig. 6.**
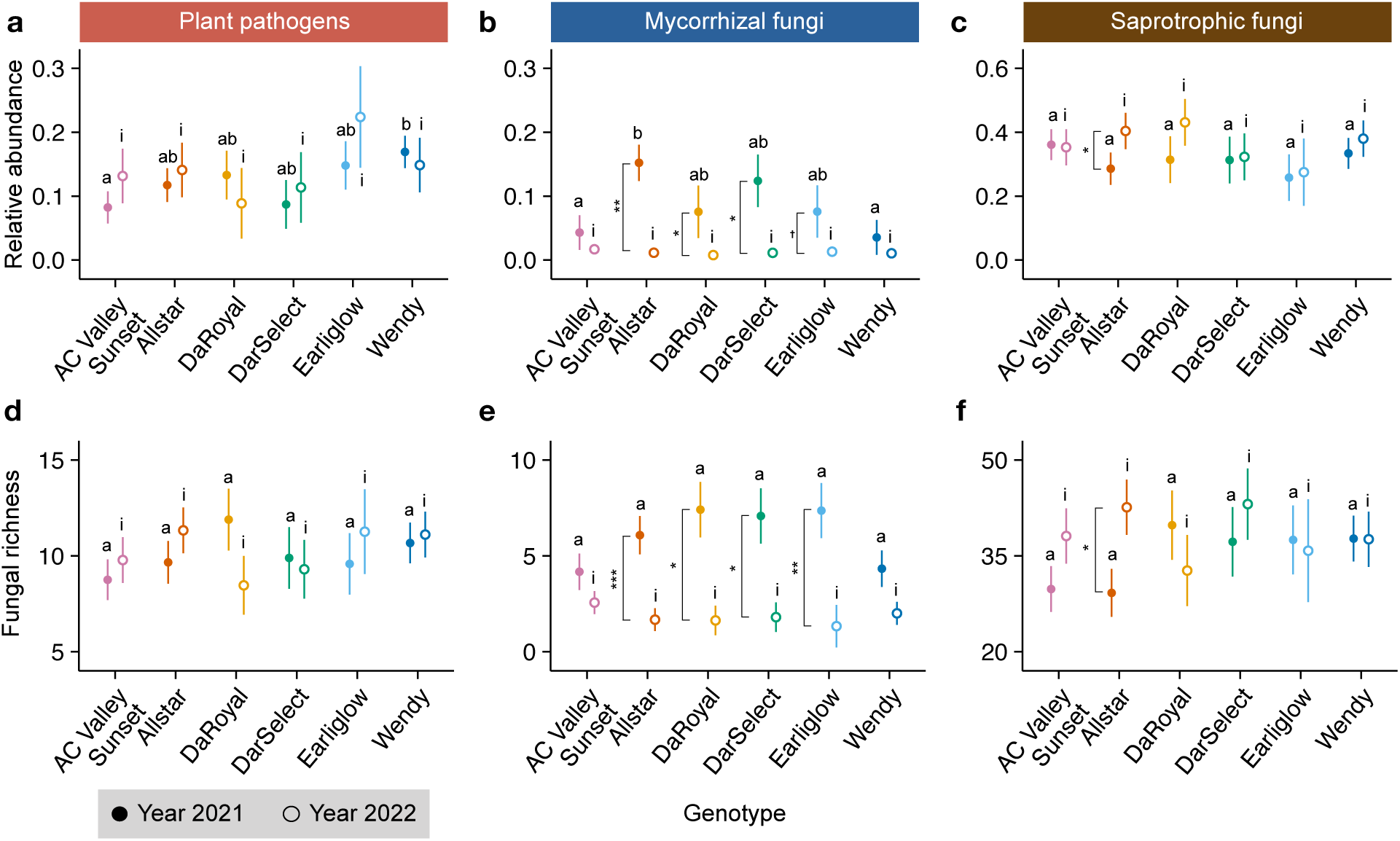
Strawberry genotypes are similar in soil fungal functional groups. The least-squares means (LS-means) of the relative abundance (a-c) and richness (d-f) for plant pathogens, mycorrhizal fungi, and saprotrophic fungi are plotted with error bars (1 SE) across, after controlling for the effect of field in general linear models (LMs) for the years 2021 (solid symbols) and 2022 (open symbols). LS-mean contrasts among genotypes are indicated by letters (a and b for 2021; i for 2022), and LS-mean contrasts between 2021 and 2022 are shown vertically. \*\*\**P* < 0.001; \*\**P* < 0.01; \**P* < 0.05.

## Discussion

This study elucidates the dynamics of soil fungal functional groups in the plant rhizosphere across fields of varying ages and multiple genotypes over time. Our results revealed an increase in the relative abundance of plant pathogens and a decrease in mycorrhizal fungi in older fields compared to younger fields, consistent with the predictions of pathogen accumulation and mutualism decline. Additionally, the relative abundance of plant pathogens increased from 2021 to 2022 in younger fields, whereas mycorrhizal fungi decreased from 2021 to 2022 consistently across fields. In contrast, the relative abundance of saprotrophic fungi remained stable across fields in both years. Despite similarities in the richness across fields in both plant pathogens and saprotrophic fungi, the community structure of both groups differed between younger and older fields. While the communities of mycorrhizal fungi were similar across fields, there was a reduction in both mycorrhizal richness and associations in older fields. The dynamics of these fungal functional groups led to variation in microbial networks across fields.

The increase in plant pathogens in older fields aligns with the prediction of pathogen accumulation over time, especially in monocultures or managed ecosystems with continuous cropping (Cook 2006; Li et al. 2014; Peralta et al. 2018; Wang et al. 2023a). The relative abundance of plant pathogens reached approximately 10–20%, which is higher than the 2–8% observed in wild grassland plants (Semchenko et al. 2018). Pathogen accumulation over time has also been reported in the experimental cultivation of wild grassland plants over four years (Kohout et al. 2024). Different from pathogen accumulation, a decline in the relative abundance of pathogens has been found during forest succession, where pathogens accounted for less than 1% across early, mid and late-successional stages (Zhao et al. 2023). This decline is, nevertheless, likely due to different host species at each successional stage, as host species with resource-acquisitive strategies, such as those in the early successional stage, tend to attract more pathogens (Semchenko et al. 2018). In this study, pathogens appeared to level off in older fields. However, this was unlikely due to pathogen saturation, as a higher relative abundance of pathogens was observed in the younger field A in 2022. We suspect the leveling off was likely due to fungicide applications to suppress soil-borne pathogens, especially in fields that were heavily infected. Indeed, more diseases, including leaf spots, fruit rot, and black root rot, were seen in the older fields. In the younger field A that was less infected, agrochemical intervention was less intense compared to heavily infected fields, potentially allowing pathogens to rise markedly over the two years. Notably, in field A, there was an increase in *Pilidium*, which can cause tan-brown fruit and root rot, as well as leaf spots in strawberries (Debode et al. 2011; Geng et al. 2012). In the older fields where black root rot disease intensified, *Rhizoctonia*, the causal agents, were present in the community composition of pathogens especially in the older fields, despite not the dominant pathogens. Different from relative abundance, pathogen richness did not increase in older fields. This was likely because the perennial plants and continuous cropping can promote the dominance of certain pathogens, such as *Fusarium*, the causal agents of *Fusarium* wilt in strawberries. In addition, agrochemical intervention could help reduce fungal richness. In this study, fertilization in urban farming may also contribute to pathogen accumulation, as nitrogen and phosphorous fertilization can increase soil-borne pathogens and disease severity (Veresoglou et al. 2012; Lekberg et al. 2021). The observed increase in pathogens in older fields and the changes over the two years highlight the accumulation and dynamics of soil-borne pathogens in the plant rhizosphere, which may affect host health not only through pathogenesis but also by influencing the dynamics of other fungal functional groups, as discussed below.

The decline of mycorrhizal fungi in older fields may suggest attenuated mutualisms over time. As pathogens accumulated in older fields, they may weaken host plants, reducing host carbon allocation to mycorrhizal fungi and competing with them for root space (Tedersoo et al. 2020). Consequently, with reduced mutualistic partners, plants would be more susceptible to pathogen infection, since mycorrhizal fungi can protect plants against pathogens through mechanisms such as competing with pathogens for space, modifying host defense chemistries, and improving nutrient access to offset pathogen-induced nutrient loss (Veresoglou and Rillig 2012; Tedersoo et al. 2020; Liu et al. 2024). In this study, we observed a reduction in the relative abundance, richness, and association of mycorrhizal fungi, which led to fewer positive network links between mycorrhizal fungi in older fields. Similar to our findings, a decline in the relative abundance of mycorrhizal fungi over time has also been observed in the experimental cultivation of wild legumes, such as *Trifolium pratense* and *Lathyrus pratensis* (Kohout et al. 2024).

However, both declines and increases in mycorrhizal fungi have been found during forest succession (Wang et al. 2023b; Zhang et al. 2024), depending on factors such as host diversity and nutrient limitations. The relative abundance (*ca*. 1–15%) and richness (3–10) of mycorrhizal fungi in this study were similar to those found in wild grassland species (abundance: *ca*. 2–10%; richness: 5–11) (Semchenko et al. 2018; Kohout et al. 2024) and lower than temperate forests (*ca*. 5–40%) (Wang et al. 2023b; Zhao et al. 2023). However, these estimates were higher than in some managed ecosystems such as maize planting fields, where the relative abundance was around 0.4% (Du et al. 2022). In addition to the older fields, from 2021 to 2022, the younger field A also experienced a marked reduction in mycorrhizal fungi, accompanied by a significant rise in plant pathogens. While negative interactions between mycorrhizal fungi and plant pathogens would predict negative correlations within microbial networks, such negative pathogen-mycorrhiza links were not yet common. This was likely because interactions can be diffused, involving multiple microbes that interact both directly and indirectly, which could weaken pairwise correlations. Alternatively, strong negative interactions might lead to the extinction or low abundance of certain microbes, resulting in their absence from the networks. In contrast to the negative interactions, positive pathogen-mycorrhiza links were observed. There could be several reasons. For example, mycorrhizal fungi vary in their ability to suppress pathogens. For some pathogens, mycorrhizal colonization does not reduce disease (Veresoglou and Rillig 2012), which could lead to positive correlations. Additionally, the effects of mycorrhizal fungi can be environment dependent, shifting from mutualistic to parasitic in nutrient-rich environments (Johnson 2010), which could also result in positive correlations with pathogens. Furthermore, competitions between mycorrhizal fungi such as mutualistic and less-beneficial ones (Werner and Kiers 2015), as well as negative interactions between pathogens and mycorrhizae, could lead to positive correlations with pathogens. In this study, the decline in mycorrhizal fungi over time might also be due to fertilization (e.g., nitrogen and phosphorous), which can alter plant investment on mycorrhizal symbiosis (Treseder 2004; Johnson 2010; Lekberg et al. 2021).

We did not observe an increase in saprotrophic fungi in older fields, contrary to our hypothesis (Fig. 1c). This was likely due to stable organic matter input across fields, as organic matter influences saprotrophic fungi (Ning et al. 2021). In this managed ecosystem–specifically, the perennial cropping of strawberries–the aboveground organic matter input from leaf litter and weeds was frequently removed to reduce disease development and maintain plant health. While accumulated pathogens in older fields may increase belowground organic matter input from dead roots, the belowground biomass in strawberries was considerably smaller than the aboveground biomass. Thus, it may not significantly increase the relative abundance of saprotrophic fungi, despite the positive correlations between pathogens and saprotrophic fungi in the microbial networks. Unlike organic matter, nutrient fertilization of nitrogen and phosphorus does not necessarily affect saprotrophic fungi (Lekberg et al. 2021; Ning et al. 2021). In natural ecosystems, the accumulation of organic matter during temperate forest secondary succession has been found to increase the relative abundance of saprotrophic fungi (Wang et al. 2023b). However, studies have also observed no significant change (Zhao et al. 2023) or a decline in saprotrophic fungi during forest succession (Zhang et al. 2024). Such variation might be attributed to shifts in host plant communities and increased nutrient limitations (e.g., N and P) that promote ectomycorrhizal fungi instead of saprotrophic fungi. In this study, the relative abundance of saprotrophic fungi was approximately 25–40%. This range varies considerably across different plants and ecosystems, such as less than 8% in subalpine forests (Zhao et al. 2023), 7–25% in temperate forests regenerated after clear-cutting (Wang et al. 2023b), and 5– 75% in pine forests (Zhang et al. 2024). While the relative abundance and richness of saprotrophic fungi remained relatively stable across fields, the community composition varied between younger and older fields and changed over the two years, indicating both shorter-and longer-term dynamics of saprotrophic fungal communities.

The genotypes did not differ in the relative abundance, richness, or community composition of plant pathogens, mycorrhizal fungi, and saprotrophic fungi. Although plant genotypes often differ in their associated bacterial and fungal communities (e.g., Wei et al. 2022; Wei and Tan 2023), the similarity in fungal functional groups across genotypes observed in this study may indicate functional redundancies and stable essential functions of microbiomes that mediate pathogenesis, mutualism, and decomposition across genotypes. Additionally, the genotypes showed similar annual dynamics of fungal functional groups, particularly a significant decline in mycorrhizal fungi over the two years. Yet, different from plant genotypes of limited genetic variation, plant species often differ in soil fungal functional groups (Semchenko et al. 2018; Zhao et al. 2023; Kohout et al. 2024). For example, the relative abundance of soil-borne pathogens can reach 40% in legumes, compared to 5–10% in other grassland species, potentially contributing to their high population fluctuations (Kohout et al. 2024). These studies and ours suggest host plants may play an important role in shaping soil fungal functional groups at broader genetic scales than genotypes.

## Conclusion

Our findings reveal contrasting dynamics among plant pathogens, mycorrhizal fungi, and saprotrophic fungi within rhizosphere soil microbiomes, which can impact plant pathogenesis, mutualism, and decomposition. As our study focused on a single ‘natural’ experiment in urban farming, it awaits similar studies to harness urban farming across cities and countries, which represents highly replicated experiments for understanding soil microbiome dynamics and functioning. Together, these studies will provide valuable insights into the repeatability and mechanisms governing the dynamics of soil fungal functional groups in the plant rhizosphere and their impact on plant productivity.

## Acknowledgements.

We thank Jessica LaBella, Elizabeth Esser, and Maris Hollowell for the assistance in field collection and microbiome DNA extraction. We also thank Bill Patterson, Dave Patterson, and other staff at the Patterson Farm for logistic support.

## Author contributions

Na Wei conceived the study, collected data, and conducted data analysis. Na Wei and Madelynn Nakaji-Conley performed data visualization. Na Wei wrote the manuscript. All authors read and approved the final manuscript.

## Funding

This work was supported by the Holden Arboretum (030869 to Na Wei).

## Data Availability

All data that support the findings of this study are included in this published article, its electronic supplementary material, and are available on Mendeley Data (DOI: 10.17632/mhvzzh6ndv.1).

## Competing Interests

The authors have no relevant financial or non-financial interests to disclose.

## Supplementary Information

The online version contains supplementary material.

## Supplementary Information

**Fig. S1.**
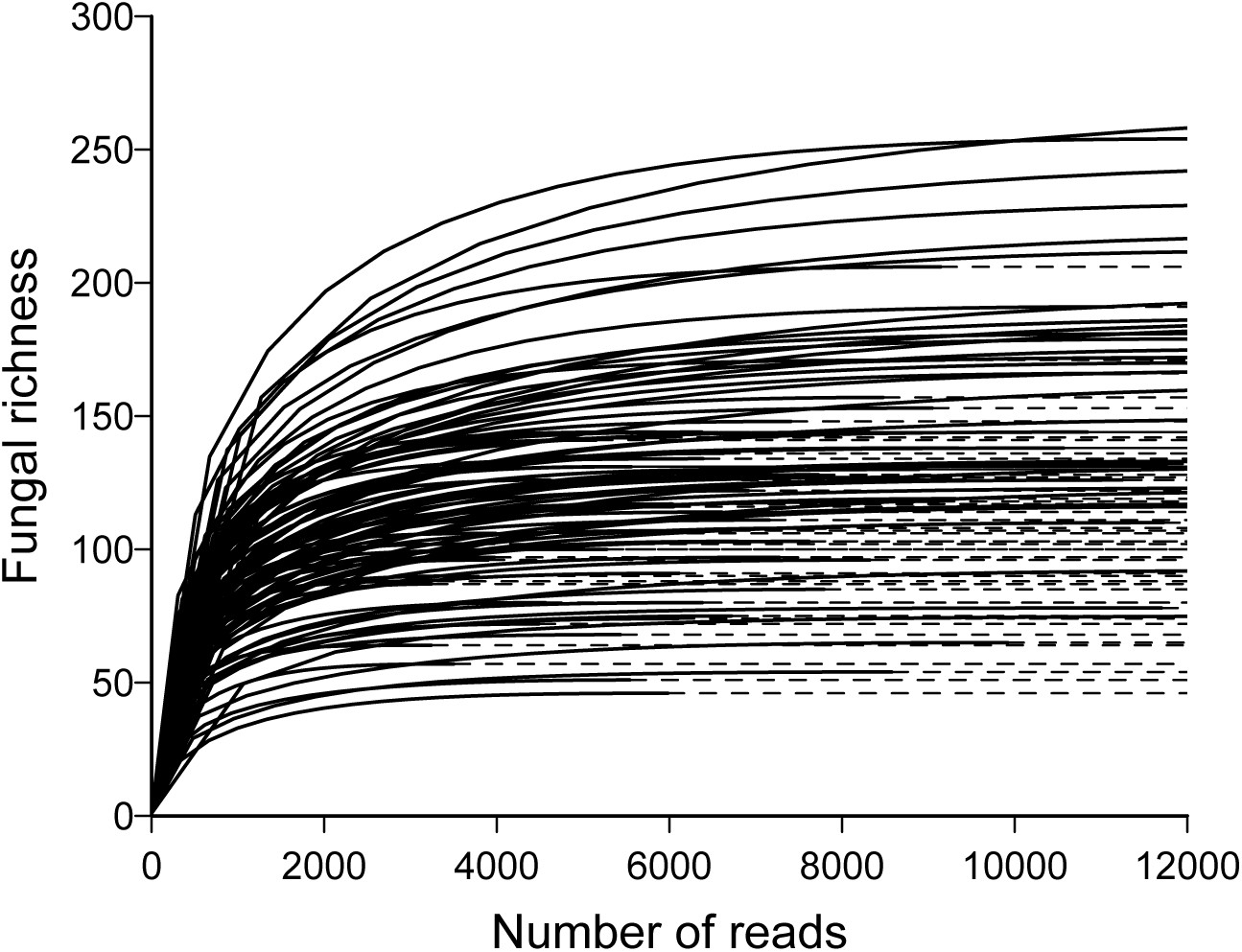
Rarefaction shows that the sequencing effort captured the majority of the fungal richness. The number of sequencing reads (*x*-axis) each sample is represented by the solid portion of each line, whereas the dashed portion indicates extrapolation in the rarefaction analysis using the R package iNEXT.

**Table S1.**
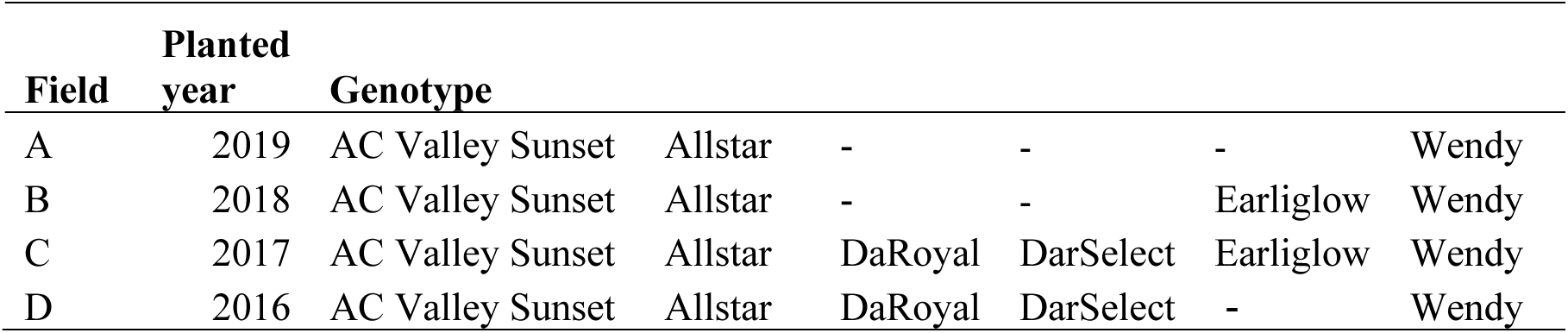
Study site information

**Table S2.** Fungal functional groups

| **Functional group** | **Number**<br>**of ASVs** | **% of**<br>**ASVs** |
| --- | --- | --- |
| Plant pathogens | 108 | 5% |
| Saprotrophic fungi | 644 | 29% |
| Mycorrhizal fungi | 136 | 6% |
| Plant endophytes | 24 | 1% |
| Parasites | 99 | 4% |
| Animal endosymbionts | 1 | 0% |
| Unassigned | 1227 | 55% |
| Total ASVs | 2239 |  |

## Notes

### Competing Interest Statement

The authors have declared no competing interest.

